# Too attractive to self: How pollinators can interfere with the evolution of selfing

**DOI:** 10.1101/2020.05.21.108225

**Authors:** RB Spigler, LM Smith-Ramesh, S Kalisz

## Abstract

Pollinators are widely invoked to explain the evolution of selfing despite genetic conditions favoring outcrossing. But their role in maintaining outcrossing despite genetic conditions favoring selfing remains unexplored. We use consumer-resource models to explicitly consider the how the plant-pollinator mutualism can constrain the evolution of selfing. We model outcrossing as a function of attractiveness and account for the cost of attractiveness as a saturating, linear, or exponential function alongside the costs of selfing: inbreeding depression and pollen discounting. We show specific, clear combinations of ecological and genetic conditions where pure selfing can invade a resident population of partial selfers. Complete selfing can evolve in the face of pollen discounting so long as there is a cost to pollinator attraction and reward. However, we also predict conditions under which mixed mating is maintained even when inbreeding depression is low. Our model highlights how under some scenarios mixed mating represents the worst of both worlds, leaving plants to pay the costs of both inbreeding depression and attraction and even leading to extinction. By linking pollinator attraction to the selfing rate, our models provide a likely common mechanism to explain pollen discounting and an alternative evolutionary pathway to the selfing syndrome.

## INTRODUCTION

The evolutionary transition from outcrossing to selfing is considered among the most common in Angiosperms (Stebbins, 1974). Classic models predicting this transition hinge on the balance between two genetic factors: inbreeding depression and an automatic transmission advantage (or cost of outcrossing) (Fisher, 1941; Nagylaki, 1976; Lloyd, 1979; Charlesworth, 1980; Lande and Schemske, 1985). Because complete outcrossers have only two pathways to pass on their alleles (i.e. outcross seed, outcross siring) but partial selfers have three (i.e. outcross seed, selfed seed, outcross siring), partial selfers should establish and spread within an outcrossing population. That is, unless the reduction in fitness of selfed offspring relative to outcrossed offspring through inbreeding depression is great enough to eliminate the 50% automatic transmission advantage of selfing. Consequently, there should be disruptive selection on the mating system. Despite early empirical data supporting this prediction (Schemske and Lande, 1985), more recent analyses indicate that 63% of species have at least one population with a mixture of selfing and outcrossing (‘mixed mating’; Whitehead *et al.,* 2018) and 12% are outcrossing despite low inbreeding depression (Winn *et al.,* 2011). Thus, the quest to understand the drivers of mating system evolution endures.

While original models emphasize the role of genetic factors, mating in most flowering plants is an ecological process involving interactions with pollinators. Not surprisingly, consideration of pollination conditions, most notably pollen limitation, has solved part of the problem of mixed mating (reviewed in Goodwillie, *et al.,* 2005; Knight *et al.,* 2005). Theoretical and empirical work illustrates how partial selfing can be favored in the face of high inbreeding depression if such selfing boosts seed production compared to outcrossed individuals when mates or pollinators are limiting (Lande and Schemske, 1985; Lloyd, 1992; Jarne and Charlesworth, 1993; Kalisz, *et al.,* 2004; Eckert, *et al.,* 2006). Indeed, selection for reproductive assurance in the face of pollen limitation is the most well accepted force favoring the evolution of selfing more generally (Busch and Delph, 2012). However, reproductive assurance alone cannot explain why outcrossing persists in spite of low inbreeding depression, underscoring that we have not yet fully explored the conditions that favor or constrain the evolution of selfing.

Explicit consideration of pollination as a mutualism involving plant and pollinator as equally interdependent actors can highlight how pollinators might constrain the evolution of selfing (Devaux *et al.,* 2014; Lepers, *et al.,* 2014; Spigler and Kalisz, 2017). Pollinators depend critically on floral resources for their own metabolic demands and for provisioning their broods. As long as this dependence exists and their local abundances are great enough, pollinators could enforce outcrossing, even if plants are capable of autonomous selfing and even if genetic conditions favor selfing (i.e., inbreeding depression is low) (Spigler and Kalisz, 2017). Near exclusive focus on the role of pollen limitation on mating system evolution has neglected this possibility, and it is worth noting that pollen limitation is often negligible and not necessarily ubiquitous (Knight *et al.,* 2005; Rosenheim *et al.,* 2014; Rosenheim *et al.,* 2016; but see Burd, 2016). Holsinger (1991) recognized the impact of pollinator abundance and plant density on the evolution of selfing and rooted his single-locus model of the evolution of mixed mating on a simple fact: the outcrossing rate is a function of how much outcross pollen is received. Also explicit is the ecological trade-off of pollen discounting: pollen used for selfing cannot be used for outcrossing and vice versa (Nagylaki, 1976b Charlesworth, 1980; Holsinger, *et al.,* 1984; Holsinger, 1991). In considering the ecological dynamics of pollinators and pollen discounting, Holsinger (1991) demonstrated theoretically not only that outcrossing could be favored in the absence of inbreeding depression but also that complete selfing will never be stable unless there is pollen limitation. We stress that this single-locus model does not account for the feedback between plant and pollinator populations that is inherent to the mutualism. As an alternative, consumer-resource models (e.g., Holland and DeAngelis, 2010) allow us to ask how pollinators can influence the evolution of selfing by explicitly connecting plant and pollinator dynamics (e.g., Lepers, *et al.,* 2014).

The economics of participating in the mutualism could also play a pivotal role in the evolution of selfing. Specifically, plants pay a price for pollinator services through floral attraction and rewards. Empirical studies of floral construction and maintenance costs of highly outcrossing flowers indicate that they can be substantial (Schemske, 1978; Waller, 1979), comprising a large fraction of a plant’s carbon budget and exacting tolls via transpiration and respiration (Nobel, 1977; Ashman and Baker, 1992; Ashman and Schoen, 1997; Ashman and Schoen, 1994; Galen, 1999; Teixido and Valladares, 2014). Although floral investment costs are not often considered in models of mating system evolution (but see Sakai, 1995 and Lepers, *et al.,* 2014), the potential for links between investment in attractiveness and the mating system is clear. Attraction should be positively correlated both with investment costs and outcrossing. These associations tack on an additional cost of outcrossing and create a clear mechanism for pollen discounting. Unattractive individuals may escape pollinators and thus achieve high selfing even in the face of high pollinator abundance, but this comes at the expense of exporting pollen and siring outcross offspring. Given these links, the question about the evolution of complete selfing becomes one about whether reductions in investment costs could enable plants to not only realize higher selfing rates but also to recoup some or all of the costs paid through inbreeding depression and pollen discounting. Indeed, across angiosperms smaller flowers are associated with higher rates of selfing (‘selfing syndrome’; Sicard and Lenhard, 2011). Moreover, as the population on average becomes less attractive and provides fewer rewards, pollinator densities could decline, which in turn would further select for increased selfing rates via reproductive assurance. We are aware of only one model that explicitly considered plant pollinator dynamics in concert with floral costs (Lepers, *et al.,* 2014), but it does not address pollen discounting.

In this study, we present consumer-resource models that consider plant and pollinator densities, selfing rate, inbreeding depression, plant attractiveness to pollinators, and the cost of attraction (or unattractiveness) including pollen discounting to evaluate the conditions under which higher rates of selfing can evolve. In contrast to most models that begin with a population of outcrossers and evaluate invasion of partial selfers, we consider whether and how higher rates of selfing can evolve within a population that is already partially selfing (mixed mating). First, we ask under what general conditions can an unattractive, completely selfing mutant replace a resident population of partial selfers whose selfing rate is a continuous function of attractiveness (Model 1). Next, we explore conditions when a less attractive, potentially more highly selfing mutant can invade a resident population of partial selfers at equilibrium with their pollinator partner (Model 2). We highlight ecological conditions that may restrict the evolution of complete selfing and uncover conditions where complete selfing may still be expected to evolve.

## METHODS

Our two models explore the relationship between pollinator dynamics and the relative success of an attractive resident vs. a less attractive invader with a higher effective selfing rate. Each of these models is based on a system of three ordinary differential equations (ODE), described below. All parameter definitions and default values are shown in Table 1. Explanations for these values can be found in Appendix A.

### Model 1: Complete vs. partial selfers

Model 1 explores the dynamics of our system where an attractive partial-selfer (*P*) competes with an invader (*5*) that exclusively reproduces through selfing. In this system, the animal pollinator *(A)* only interacts with the partial selfer, although it can maintain a positive population size as a ‘generalist’ feeding on other flowers that are external to the two-species system.

The growth rate of the resident partial selfer’s population density, *P,* is given by equation 1. The resident produces *r* ovules per individual per unit time, representing its maximum intrinsic growth rate. These ovules can either be fertilized through outcrossing or selfing, where outcrossing occurs according to the rate of interactions between plants and pollinators. Pollinators visit flowers according to their attractiveness, *a_P_*, as a saturating function of plant and animal densities with a half-saturation constant *h_1_*. Any ovules that are not fertilized through outcrossing are fertilized through delayed selfing (we assume no self-pollen limitation), with survival discounted by inbreeding depression, *δ*. The resident plant experiences density dependent mortality according to the density of *P* and *S* individuals combined, which are assumed to be competing for common resources. The resident also experiences losses through the cost it pays to produce attractive flowers and nectar for pollinators, where cost is a saturating function of attractiveness, *a_P_*, with a half saturation constant *h_2_*, up to a maximum per capita cost *c.* The cost of attraction as a single metric is supported by empirical evidence of positive correlations between flower size and reward in some systems (e.g., Stanton and Young, 1994; Campbell, 1996; Fenster *et al.,* 2006; Tavares, *et al.,* 2016).

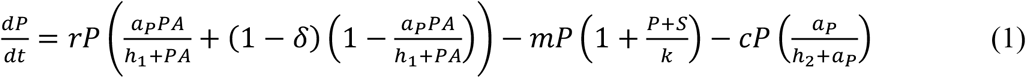

The invading complete selfer population density, *S,* also produces *r* ovules per individual, and since all ovules are fertilized through selfing, successful fertilization is discounted by inbreeding depression, *δ.* Because the selfer is completely unattractive to pollinators, there are no mating events between the complete selfers and the resident partial selfers. The invader experiences the same density-dependent mortality as the resident. Its growth rate is given as:

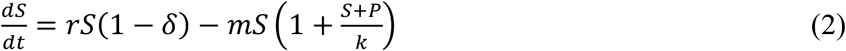

The animal pollinator population density, *A,* grows based on its consumption of floral resources, represented by an intrinsic growth rate *ρ* that is independent of the resident plant density. The pollinator population can also benefit from feeding on the resident partial-selfer, where *β* is the maximal per-capita benefit and the actual benefit is a saturating function of plant density. The pollinator also experiences density-dependent mortality, with a maximum mortality rate of *μ* scaled by a density-dependent factor *K.* Its growth rate is defined as:

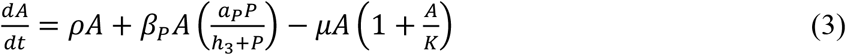

The model was run according to default parameter values shown in Table 1.

### Model 2: Invasion dynamics of a less attractive mutant

Model 2 explores a case wherein a mutant genotype *(M)* arises in a resident population of partial-selfing plants (*P*) fertilized by a generalist animal pollinator (*A*). The mutant genotype is less attractive than the resident (*a_M_* < *a_P_*) but is otherwise biologically identical. Because the mutant arises from the resident population, it is initially very rare (one individual) and arises in a resident population that is initially at equilibrium with its animal pollinator. Consistent with previously validated phenotypic models (Cheptou, 2004; Lepers, *et al.,* 2014), we assume that when outcrossing occurs between resident and mutant genotypes, 50% of the offspring exhibit the resident phenotype, while 50% exhibit the mutant phenotype.

This model consists of an ODE with three equations, one for the resident partial-selfer (*P*), one for the less attractive mutant (*M*), and one for the animal pollinator (*A*).

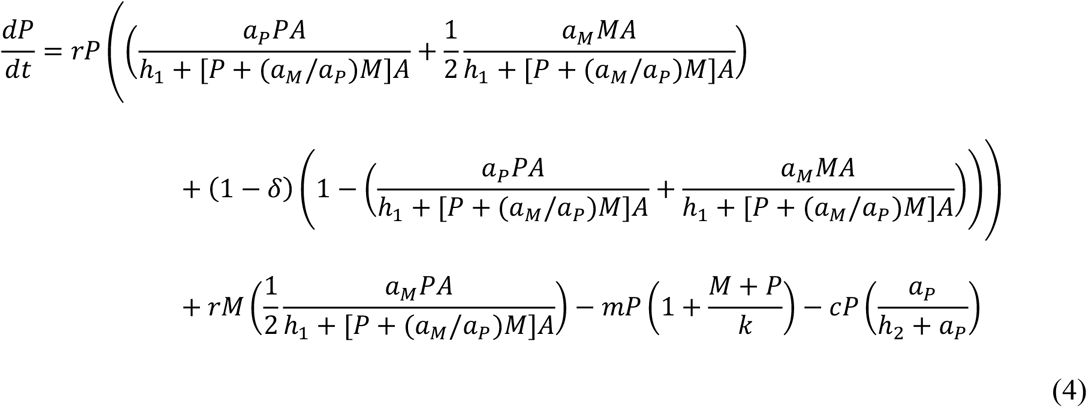

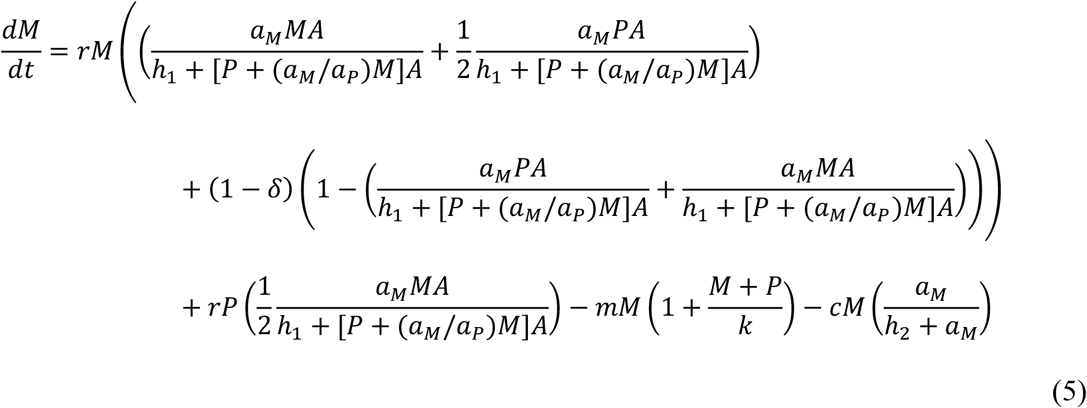

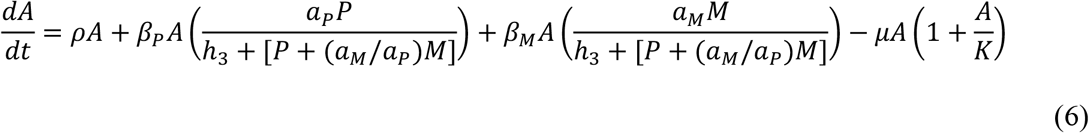

The resident partial selfer population density, *P,* grows according to the fraction of its total ovule production (*rP*) devoted to outcrossing with other residents, outcrossing with mutants, and selfing (Eqn 4). Within the parentheses following *rP*, the first term indicates the fraction of ovules fertilized by the pollinator with pollen from other plants with the resident genotype. Outcrossing is a saturating functional response (half saturation constant *h_1_*) of plant and pollinator densities, and depends upon the attractiveness, *a_P_*, of the resident plant, with 100% of these ovules result in an offspring with the resident phenotype. The second term indicates the fraction of ovules fertilized by the pollinator with pollen from plants with the mutant genotype. Again, outcrossing is a saturating functional response of plant and pollinator densities but depends upon the attractiveness of the less-attractive mutant, *a_M_*, such that when the mutant is very unattractive, between-genotype outcrossing occurs at a low rate regardless of the attractiveness of the resident, *a_P_*. From these ovules, 50% of offspring exhibit the resident phenotype, while 50% exhibit the mutant phenotype. The remaining term within the parentheses following *rP* indicates the fraction of ovules produced by the resident that are not fertilized by the pollinator, and therefore are fertilized through delayed selfing. Of these self-fertilized ovules, a fraction (1-δ) survives, accounting for mortality due to inbreeding depression, δ. As with Model 1, we assume no pollen limitation; all ovules not fertilized with outcrossed pollen are fertilized with self-pollen. In the next model term, *rM* indicates total mutant ovule production, and the term following *rM* indicates the fraction of these ovules fertilized through outcrossing with resident plants that result in a resident phenotype. Finally, the resident plant experiences losses through density dependent mortality according to a maximum mortality rate *m* and a density dependent term *k.* The resident also experiences a cost of pollination that is independent of pollinator density but increases as a function of attractiveness following three alternative cost functions (figure 1). In the first scenario the cost of pollination increases to a maximum cost *a_P_* at a saturating rate with attractiveness, *a_P_*, based on a half-saturation constant *h_2_* (shown in text). In the second, the cost of pollination increases linearly with attractiveness. In the third function, the cost of pollination increases at an accelerating rate as the plant becomes more attractive.

**Figure 1.**
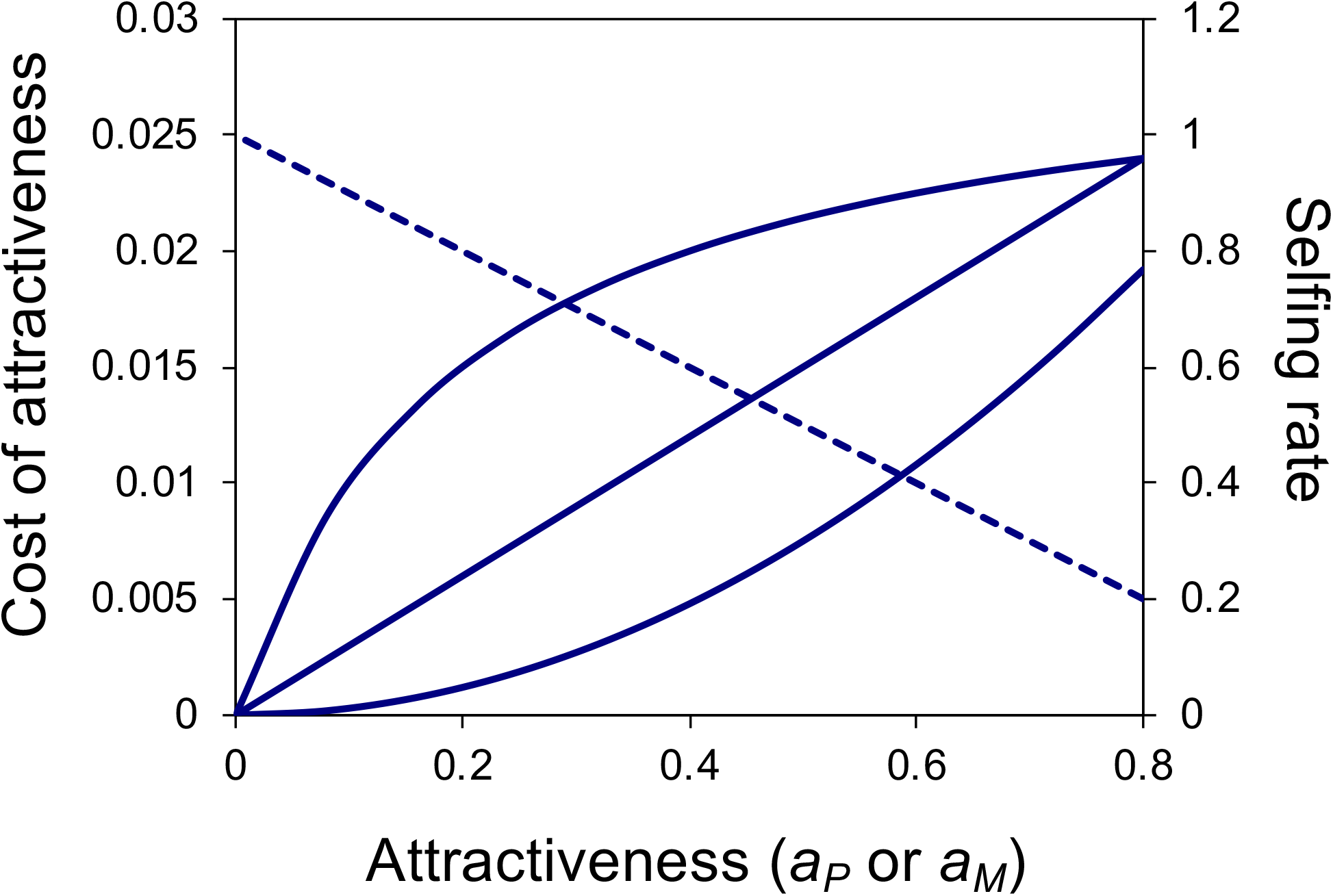
Cost and selfing rate as a function of attractiveness. We consider three cost function curves as shown (saturating, linear, accelerating) on the primary axis and indicated by the solid lines. Maximum cost value, *c*, is shown here as *c* = 0.03. Selfing rate decreases as a linear function of attractiveness, represented by the dotted line in reference to the secondary axis.

The less attractive mutant population density, *M,* grows according to an identically structured equation (Eqn 5), where the primary difference between species is that the mutant is less attractive than the resident (*a_M_* < *a_P_*).

The animal pollinator population density, *A*, grows based on its consumption of the two plant genotypes as well as through external floral resources (Eqn 6). When the pollinator is a generalist, it feeds on external floral resources, represented by an intrinsic growth rate *ρ* that is independent of the resident and mutant genotype densities. The pollinator population density can also increase through feeding on the resident partial selfer and the mutant, where *β_i_* is the maximal per-capita benefit and the actual benefit is a saturating function of genotype densities with a half-saturation constant *h3*. The pollinator also experiences density-dependent mortality, with a maximum mortality rate of *μ* scaled by a density-dependent factor *K*.

The model is written on the assumption that *a_M_* ≤ *a_P_*, and therefore conditions where a mutant that is more attractive than the resident partial selfer never exist. This assumption plays into the model structure in the attractiveness terms, where the probability of outcrossing between genotypes depends on the attractiveness of the mutant rather than the resident. This assumption also influences the denominator of the pollination functional response, where pollinator satiation based on mutant abundance is weighted by the relative attractiveness of the mutant. We note that where *a_M_* = 0 (i.e., where the attractiveness differential = 1), the mutant is a complete selfer and this model collapses to the case of Model 1.

### Model 2 Analysis

We analyzed the model to determine under which conditions the mutant (*M*) would meet the invasion criterion of exhibiting a positive population growth rate when rare when the resident partial selfer (*P*) and the animal pollinator (*A*) were at equilibrium. The two-equation system for the resident genotype and pollinator could not be solved analytically, so equilibrium densities were calculated through numerical simulations. Under each set of parameter values, the model was simulated for 5000-time steps in the software program *Mathematica,* using the ‘NDSolve’ numerical integrator. This resulted in convergence on stable densities for both the resident genotype, *P*, and its animal pollinator, *A*, under all conditions presented here. After equilibrium densities were calculated for all relevant scenarios, initial mutant growth rate was calculated based on Eqn 5.

We note that Model 2 is written to apply when the mutant is rare and that the simple relationship between genotype and phenotype that we assumed here is unlikely to apply when the mutant is common. As such, in contrast to Model 1, we cannot view Model 2 as an equilibrium model. Instead, we analyze Model 2 based on the invasion criterion of the mutant exhibiting a positive population growth rate when rare and cannot speak to the outcome of the mutant invasion in the long-term.

## RESULTS

### Model 1: Complete vs. partial selfer

Model 1 examines the outcome when a partial selfer competes with a complete selfer. Strikingly, we find a large parameter space over which the complete selfer wins (figure 2), even in the face of complete pollen discounting. When the resident is highly attractive (*a_P_* = 0.8), the complete selfer can win at low inbreeding depression levels only *(δ* <0.25). At such levels, the complete selfer fertilizes all ovules but pays neither the cost of making flowers nor a substantial cost of inbreeding. Therefore, it replaces the partial selfer, who pays a high cost of attractiveness. However, as inbreeding depression increases, the balance tips in favor of the partial selfer. Although the partial selfer still pays the same cost of making flowers, its high rate of outcrossing allows it to largely avoid the cost of inbreeding depression. The complete selfer, however, pays the cost of inbreeding depression on all progeny. The partial selfer can persist even when inbreeding depression is complete (5=1) because it maintains maximum levels of outcrossing when it is highly attractive (figure B1), though equilibrium density of the partial selfer declines with increasing inbreeding depression (figure 2 *A*).

**Figure 2.**
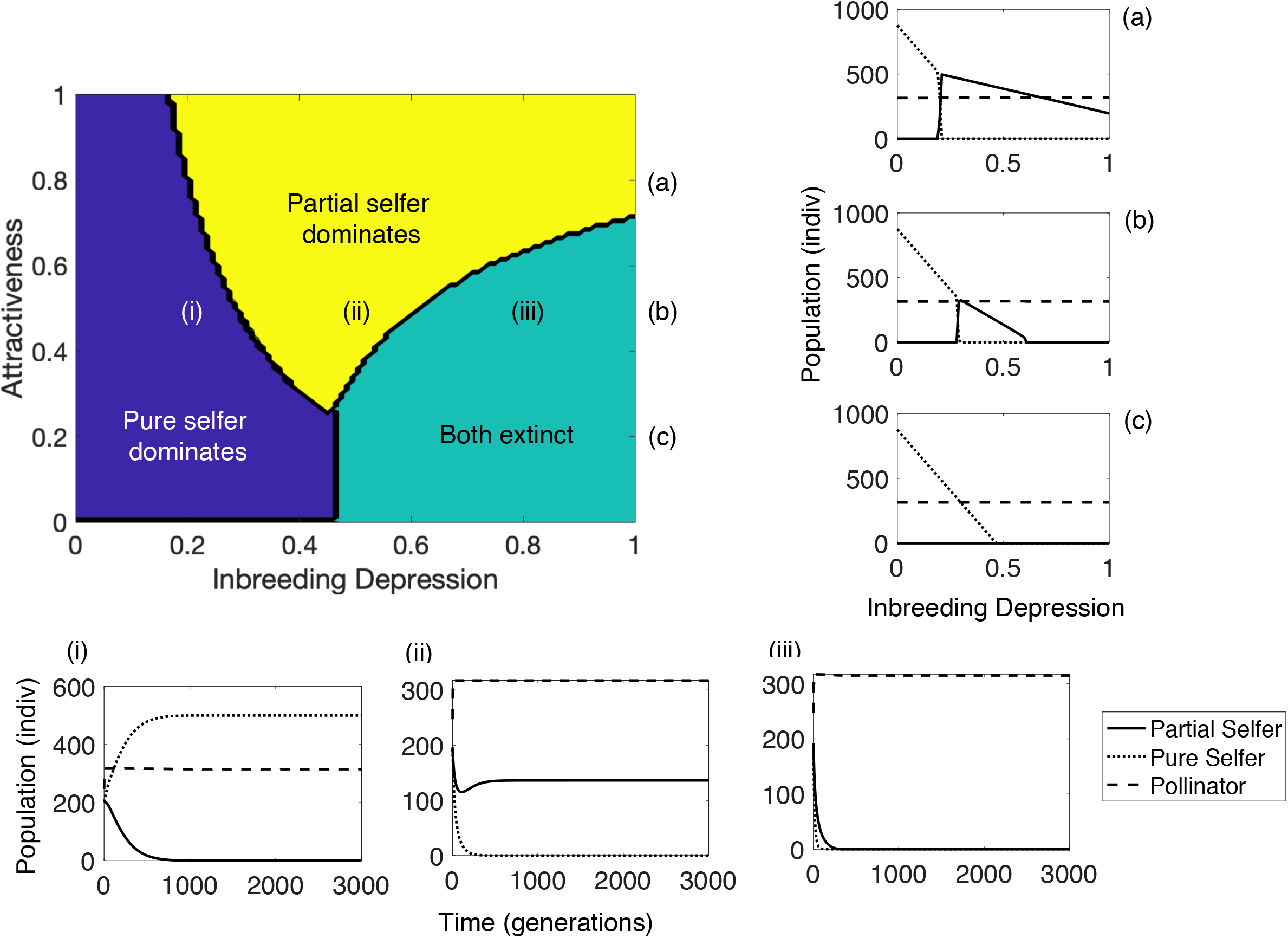
Evolutionary outcome for the evolution of complete selfing depending on inbreeding depression (*δ*) and attractiveness of a resident partial selfer (*a_p_*) that depends on a generalist pollinator. The dark blue region indicates where the pure selfer dominates, and the yellow region indicates dominance of the partial selfer; neither morph persists in the aqua region. To better illustrate the underlying population dynamics, we show population sizes of the partial selfer, pure selfer, and pollinator as a function of inbreeding depression when (a) *a_p_* = 0.8, (b) *a_p_* = 0.5, and (c) *a_p_* = 0.2. We also show how population sizes of the partial selfer, pure selfer, and pollinator change over time when *a_p_* = 0.5 and (i) *δ* = 0.2, (ii) *δ* = 0.2, and (iii) *δ* = 0.2.

As the attractiveness of the resident partial selfer decreases, the complete selfer wins at increasingly higher levels of inbreeding depression while the range over which the partial selfer can win becomes greatly restricted, instead going extinct. When the partial selfer is moderately attractive (e.g., *a_P_* = 0.5), the dynamics for the complete selfer do not change drastically from the case of the highly attractive partial selfer (complete selfer now wins up to δ ~0.3), but the partial selfer only wins when inbreeding depression is between 0.3 and 0.6 and declines to extinction when δ >0.6. The moderate reduction in attractiveness has two key consequences for the partial selfer. First, under a saturating function this reduction in attractiveness barely reduces the cost of attractiveness compared to the case of *a_P_* =0.8 (figure 1). Second, the partial selfer now has a higher selfing rate and so incurs a higher cost of inbreeding for a larger proportion of seeds. This combination represents the worst of both worlds associated with mixed mating, resulting in greater net losses for the partial selfer. Consequently, the pure selfer wins at marginally greater inbreeding depression values because it still pays no cost of attraction, and the partial selfer occurs at a lower density even where it wins (figure 2 *A* vs 2 *B*). At even lower levels of attractiveness (e.g., *a_P_* = 0.2), the partial selfer fares even worse because it now self-fertilizes at a rate nearly identically to the complete selfer yet continues to pay some cost of attractiveness. This scenario tips the scale in favor of the complete selfer, such that the complete selfer wins up to the boundary value of *δ*=0.5. After that point, neither type of selfer persists (Fig 2 *C*).

### Model 2: Invasion dynamics of a less attractive mutant

With Model 2, we fix the attractiveness level of a resident partial selfer and asked whether a less attractive mutant can invade when rare. The attractiveness of the mutant is defined relative to that of the resident in terms of an ‘attractiveness differential’. When the resident is highly attractive (*a_P_*=0.8) and the maximum cost of pollination is set at *c*=0.03, a saturating cost function means that initial decreases in the mutant’s attractiveness relative to a highly attractive resident—represented as low values of the attractiveness differential—do not appreciably decrease floral investment costs (figure 1). Yet the selfing rate still increases linearly (figure 1). This combination combined with inbreeding depression strongly restricts the ability of the mutant to invade, even at *δ*<0.1 (figure 3 *A*). Once the attractiveness differential reaches ~ 0.5 or greater, however, the savings in investment costs for the mutant increases, and the benefits of reduced flower/reward costs begin to offset the cost of greater selfing at even higher inbreeding depression levels. This benefit is reflected in the rapidly growing region in which the mutant can invade as attractiveness differential continues to increase (figure 3 *A*). Even still, the mutant overall remains limited by inbreeding depression. Note that at an attractiveness differential of 1 (*a_P_*=0.8 and *a_S_*=0), results are identical to our first model for *a_P_*=0.8, since the mutant is a complete selfer.

**Figure 3.**
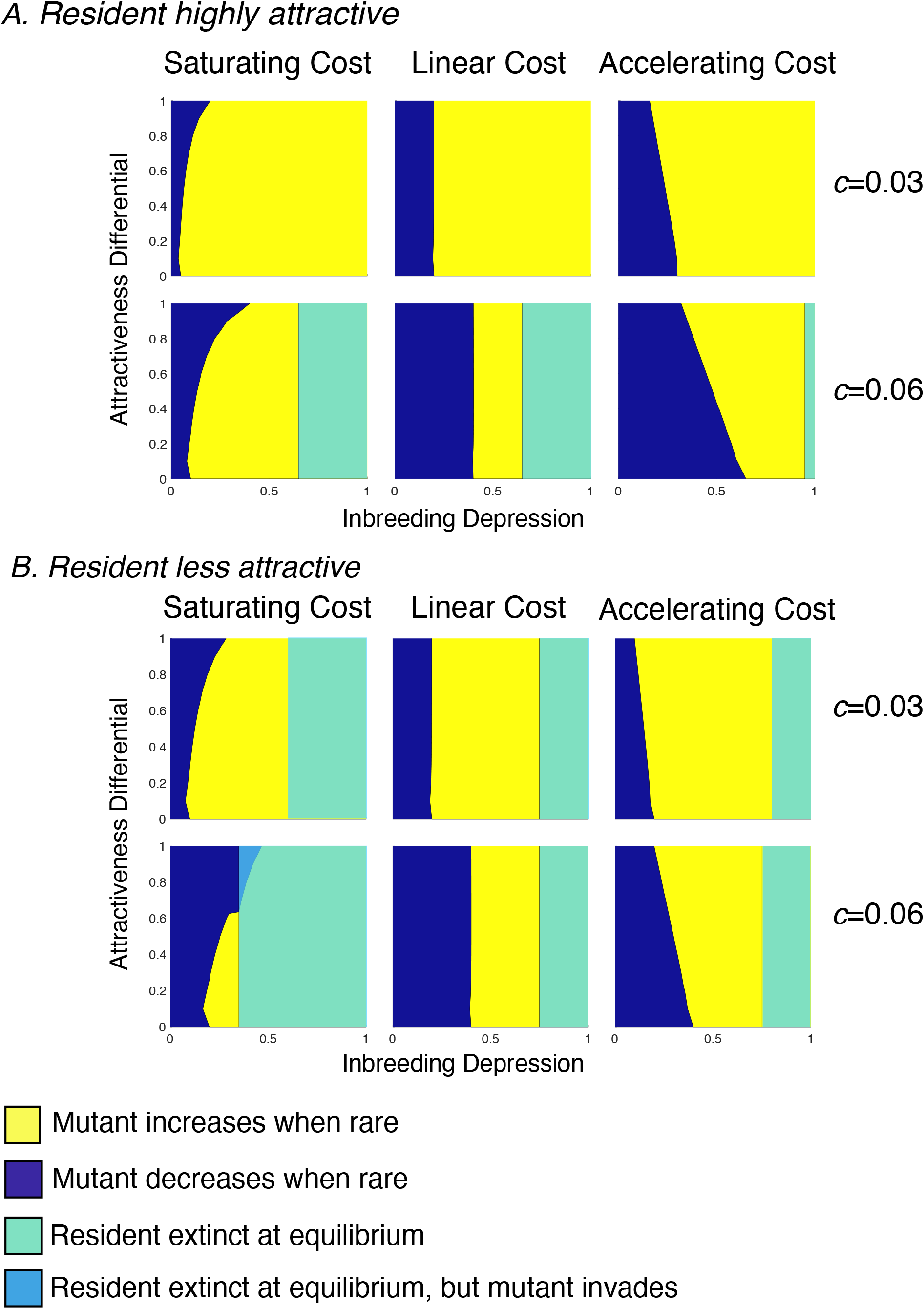
Pairwise invisibility plots for three different cost functions (saturating, linear, accelerating) and two maximum costs of attractiveness (*c*). A. Resident is highly attractive (*a_p_* = 0.8). B. Resident is less attractive (*a_p_* = 0.5). Four possible outcomes are represented according to the figure legend: the mutant can successfully invade (increases when rare), the mutant cannot invade (decreases when rare), density of the resident is 0 at demographic equilibrium, and density where the resident is 0 but the mutant can invade.

Compared to a saturating cost function, linear and exponential curves always result in a lower cost for a given level of attractiveness (figure 1) and so create more favorable conditions for a less attractive, more highly selfing mutant to invade when the resident is highly attractive (figure 3). Now, the mutant can invade over a greater combination of attractiveness differentials and inbreeding depression values than it can under a saturating curve. When the cost of attraction is an increasing linear of attractiveness, it is inversely related to the selfing rate. Consequently, any change in attractiveness (and thus cost of attraction) is balanced by a proportional change in selfing rate (and thus proportion of individuals paying the cost of inbreeding depression). This creates a critical threshold value of inbreeding depression; below this threshold the mutant can invade at any attractiveness differential (figure 3). As long as inbreeding depression is below this threshold, any losses the mutant pays from inbreeding depression will be outweighed by gains from the reduction in attraction costs, and the initial growth rate of the mutant is positive.

When instead the cost function is exponential, the mutant is more successful at invading at lower attractiveness differentials (figure 3). Under this cost function, small reductions in attraction relative to a highly attractive resident (*a_P_*=0.8) result in a dramatic decrease in costs that is disproportionate to the linear increase in selfing rate. Hence, the savings from reduced attractiveness offset the losses to inbreeding depression. With further decreases in attractiveness under and exponential function, however, the reduction in cost diminishes. Functionally, this leads to a slight retraction of the range of inbreeding depression values over which the mutant can invade as the attractiveness differential increases.

We find that the impact of altering the attractiveness of the resident depends on the shape of the cost function (figure 3 *B* vs. 3 *A*). When the resident is only moderately attractive (*a_P_*=0.5), the region over which the mutant can invade expands under a saturating cost curve, remains the same under a linear cost curve, and contracts under an exponential curve. Under a saturating curve, expansion arises when the resident starts out less attractive (*a_P_*=0.5) because the mutant’s lower attractiveness finally begins to pay off in terms of cost savings (see figure 1). The reverse occurs under an exponential cost function at *a_P_*=0.5 because the reduction in attractiveness costs with the reduction in attractiveness begins to slow down, and difference in floral costs paid by the mutant at increasingly greater attractiveness differentials becomes minimal. In addition, a less attractive resident exhibits negative growth rates and goes extinct over a range of inbreeding depression values, with the threshold dictated by the cost curve (figure 3). As in Model 1, this consequence arises because moderate attractiveness results in the worst of both worlds scenario when inbreeding depression is high.

When we double the maximum cost of pollination to the resident (*c*=0.06), the shape of the parameter space remains the same, but the size of the space in which the mutant can successfully invade doubles (figure 3). Under this scenario, the equilibrium density of the resident is lower compared to the case of *c*=0.03, making it easier for the mutant to have positive growth rates and successfully invade at higher inbreeding depression levels. Another key difference when the maximum pollination cost is greater is that the resident goes extinct over a larger range of parameter values. For a highly attractive resident (*a_P_*=0.8), we see this across all cost curves (compare top and bottom panels of figure 3 *A*), though inbreeding depression threshold above which the resident goes extinct varies across cost curves. When the resident is moderately attractive (*a_P_*=0.5), the level of inbreeding depression at which it goes extinct decreases slightly under an exponential cost function, remains the same under a linear cost function, and drastically decreases under a saturating cost function such that extinction is predicted even when *δ*<0.5.

## Discussion

With our model we considered whether the role of pollinator availability can cut both ways. That is, if a lack of pollinators can sustain mixed mating despite genetic conditions selecting against selfing, can the presence of pollinators also sustain mixed mating despite genetic conditions favoring greater selfing? If so, what are the conditions that define how and when selfing can evolve when pollinators are abundant? Our consumer-resource modeling approach reveals how the economics of floral investment, pollen discounting, and inbreeding depression interact to maintain mixed mating in species capable of autonomous selfing and finds limited circumstances under which greater selfing rates can evolve when pollinators are abundant.

### Pay to Play

Despite inclusion of pollination biology in theoretical models of mating system evolution (reviewed in Goodwillie, Kalisz and Eckert, 2005), the cost of participating in the pollination mutualism by generalist-pollinated plants has largely been ignored (but see Lepers, Dufay and Billiard, 2014; Sakai, 1995). Most mutualisms, however, incur a cost (Bronstein, 2001), including generalist pollination systems (Morris, Vázquez and Chacoff, 2010). Key to remaining in the mutualism is whether the benefits provided outweigh such costs. For self-compatible plant species, the alternative to investing in pollinator attraction is to fail to attract pollinators, self-fertilize, and pay the potentially greater toll inflicted by inbreeding depression. Thus, to outcross, plants must pay to play. When the cost of attraction enters the mating-system evolution equation, the situation becomes about whether reductions in investment costs enable plants to both realize higher selfing rates and recoup the costs paid through inbreeding depression. Not surprisingly then, we find that increasing maximum floral investment cost increases the parameter space over which more highly selfing individuals can invade.

Consideration of the level of attractiveness of the resident partial selfer and its interaction with the shape of the associated cost function leads to new and surprising results. In our first model considering a pure selfer and a saturating cost curve, for example, we show that it is more difficult for a complete selfer to win against a *more* attractive, partial selfer than a *less* attractive partial selfer. This result is somewhat counterintuitive, since the greater the attraction, the greater the floral costs paid by the partial selfer. But, attractive plants outcross and in doing so avoid paying the cost of inbreeding depression. For a completely attractive plant in Model 1, paying the cost of attraction alone is less expensive than the cost paid by a pure selfer in the currency of inbreeding depression, until inbreeding depression is low enough such that the balance is tipped in favor of the pure selfer (< 0.2 based on the parameter values modeled here). In contrast, less attractive partial selfers begin paying the cost of inbreeding depression on top of floral costs, paving the way for pure selfers to win at increasingly greater levels of inbreeding depression. This result highlights the fact that mixed maters can get stuck paying double, and when the combined costs of inbreeding depression and attraction become too great, go extinct. Nevertheless, these dynamics create a parameter space over which mixed mating (0.2<s<0.8) and even outcrossing (s<0.2) can win against a pure selfer when ID<0.5.

When we consider different cost functions and investigate conditions under which higher selfing rates may evolve via the invasion of a less attractive mutant, we find that the shape of the cost function can drive mating system evolution and the potential for the stability of mixed mating. In general, the evolution of selfing is more permissive under linear and exponential cost functions than under a saturating one. However, we reveal an interaction between the resident’s attractiveness and the shape of the cost function: as the attractiveness of the resident increases, the evolution of higher selfing rates becomes more permissive under an exponential cost function but less permissive when the cost curve is saturating. Although there exist limited cost estimates related to floral construction or maintenance (e.g., Oakley, Moriuchi and Winn, 2007; Ashman and Schoen, 1994; Ashman and Schoen, 1997; Galen, 1999), the shape of the cost function is not known for any species. Our model results suggest that variation among species in cost functions, due to differences in factors such as reward type, flower size and floral longevity, could drive variation in the conditions necessary for higher selfing rates to evolve.

Our approach complements and extends the work by Lepers et al. (2014). They too emphasized the co-evolution of mating system and floral traits and showed how, in some cases, pollinators may interfere with the evolution of selfing. However, there are important differences between the studies. First, Lepers at al. (2014) considered a broader set of conditions including a gradient of pollinator specialization. When the mutualism is highly specialized, plant and pollinator densities are tightly coupled, which can feed back to create simultaneous mate and pollinator limitation and favor selfing. In this way, Lepers at al. (2014) more fully take advantage of consumer-resource dynamic feedbacks. However, we restricted our analysis to the case of the more common generalized pollinator (Waser *et al.,* 1996; Ollerton, 1996; Thomson, 2003) because our goal was to explore whether and how selfing could arise without pollinator limitation. Second, and to that end, although both studies consider autonomous selfing, we elected to model delayed selfing, while Lepers et al. (2014) modeled prior selfing (sensu Lloyd, 1979). Prior selfing could provide another avenue by which plants may be able to evolve selfing in the presence of pollinators (Brys, *et al.,* 2016; Randle, *et al.,* 2018; Spigler and Kalisz, 2017) as could reduced herkogamy, but our emphasis on delayed selfing further allows for the elimination of pollen limitation as plants compensate for reductions in outcross pollen receipt (i.e. reproductive assurance). In addition, because the correlation between selfing rate and attractiveness in our model arises from pollinator behavior and not underlying genetic architecture, we create a scenario where pollinators enforce outcrossing, flipping the sign of the interaction to negative for the plant when inbreeding depression is low. Delayed selfing with low inbreeding depression might not be that uncommon (Goodwillie and Weber, 2018). Finally, the model of Lepers et al. (2014) is free from pollen discounting, the reduction in pollen export with increased self-pollination, a potentially critical parameter shaping mating system evolution (Nagylaki, 1976; Charlesworth, 1980; Holsinger, Feldman and Christiansen, 1984; Johnston *et al.,* 2009), whereas it an essential component in ours. Ultimately, while our model may represent a more limited case, we are uniquely able to account for the presence of mixed-mating species with low inbreeding depression (Winn *et al.,* 2011).

### Pollen discounting

In our models, both the resident and less attractive mutant can experience pollen discounting as a function of their attractiveness. Unattractive flowers self-pollinate more because they receive fewer pollinator visits; yet because they do not receive as many visits they also export less. In modeling this connection, we gain another novel outcome: even with pollen discounting and abundant pollinators, the evolution of complete selfing is possible. This occurs because less attractive, more highly selfing individuals are able to recoup or even overcome the costs of inbreeding depression and pollen discounting through reduced floral expenses. Pollen discounting in our model is independent of the cost of attractiveness, but an emergent property is that the severity of pollen discounting increases non-linearly with inbreeding depression (figure B2). This is because as inbreeding depression increases, the *gains* to the mutant via selfing decrease by a constant percentage while siring *gains* remain the same. This could also be explained by density since resident density declines linearly with inbreeding depression (figure B3). We see that selfing *rate* is impacted neither by inbreeding depression nor density of the resident, but siring *rate* increases in proportion to density with higher rates of inbreeding depression. We cannot say the degree to which pollen discounting and the change in its severity with inbreeding depression (or density) influences our results relative the influence of the cost of attractiveness. In particular, because the mutant is so rare, its highest siring rate is <1% of resident ovules when resident density > 0. Nevertheless, these patterns highlight the importance of considering the links between pollinator visitation, plant density, inbreeding depression, and pollen discounting and suggests that selfers could pay an even higher cost of selfing when inbreeding depression is high. Future modeling efforts will explicitly account for the role of pollen discounting and its context dependency.

### Caveats

Several features of our model and assumptions may limit its generality. Although consumer-resource dynamic modeling can expand our understanding of the roles that demography and ecological interactions have in mating system evolution, they are inherently phenotypic, demographic models, not genetic models. Therefore, we can only count ovules produced and cannot properly account for the transmission advantage, nor how it may change as the frequency of selfing increases in the population (Holsinger, 1991). In addition, we note that inbreeding depression stays constant within our model. Inbreeding depression may evolve alongside the selfing rate and so influence the outcome of mating system evolution (Lande and Schemske, 1985; Charlesworth and Charlesworth, 1987; Charlesworth and Charlesworth, 1990; Charlesworth, *et al.,* 1990; Husband and Schemske, 1996; but see Lande, *et al.,* 1994 and Winn *et al.,* 2011). However, because our study is primarily concerned with initial invasion conditions over relatively short ecological time periods, the assumption of equivalent inbreeding depression levels between the mutant and resident is reasonable. Inclusion of additional parameters or correlations such as those between flower size and ovule number or flower size and number (e.g., Worley and Barrett, 2000; Worley and Barrett, 2001; Caruso, 2004; Delph *et al.,* 2004; Spigler and Woodard, 2019) would undoubtedly influence our outcomes. Further, pollinator-mediated selfing is influenced by floral display size and flower size and can lead to pollen discounting (Harder and Barrett, 1995) but is not consider here. Finally, we assumed that all genotypes are equally capable of autonomous selfing, such that selfing ability, per se, and floral attraction (such as flower size) vary independently. Future models can investigate variation in both attraction and selfing ability to examine their joint evolution and outcomes for the evolution of selfing and the selfing syndrome.

### Evolutionary implications

Our model provides an alternative hypothesis for the origin of the selfing syndrome. We consider: what if small flower size is what allows plants to achieve higher selfing rates to begin with? That is, if pollinators are abundant and enforce outcrossing, then high rates of selfing can only be achieved if flowers are unattractive. Our results illustrate how pure selfers only have an advantage if their floral investment is sufficiently reduced. We recognize that because we are modeling a case without pollen limitation, where individuals are already capable of autonomous selfing, that this scenario may be applicable under a restrictive set of cases. Nevertheless, it provides an alternative pathway to the common association between flower size and mating system in angiosperms (Sicard and Lenhard, 2011).

Our models also lead to predictions about the success of mutations with varying effect sizes. Understanding the genetic basis of adaptation and the distribution of underlying effect sizes represents active areas of theoretical and empirical research (Yeaman and Whitlock, 2011; Savolainen, *et al.,* 2013; Dittmar *et al.,* 2016). Studies of floral and mating system traits have found evidence for both alleles of large and small effects (Bradshaw *et al.,* 1995; Bernacchi and Tanksley, 1997; Fishman, *et al.,* 2002; Goodwillie, *et al.,* 2006; Slotte *et al.,* 2012; Ferris *et al.,* 2017). Our models investigating the invasion of a more highly selfing mutant suggest that the success of large vs. small effect alleles is highly context dependent, determined by the shape of the cost function, maximum floral cost, attractiveness of the resident, and inbreeding depression. Under a saturating cost function, a mutation of large effect, creating a large attractiveness differential between the resident and mutant, will be successful at invading over a wider range of inbreeding depression values and may allow for a more rapid evolutionary shift to higher selfing. In contrast, invasion success of alleles of small effect is highly restricted to only the lowest inbreeding depression levels. Lower maximum floral costs also translate into the need for a much larger effect mutation for higher selfing to evolve for a given inbreeding depression level under a saturating curve. For example, under conditions of *δ* = 0.25 and *ap=0.5* (figure 3 *B*), we find that a more highly selfing mutant can successfully invade when c=0.06 so long as the mutation(s) results in >50% reduction in attraction relative to the resident. But, when *c*=0.03 only a mutation resulting in a > 90% reduction—near complete selfing—would be successful. A similar effect occurs when we hold the maximum floral cost and inbreeding depression constant for a saturating curve but alter the attractiveness of the resident; the threshold for a small effect mutation to invade is much lower when the resident is less attractive. For a linear cost function, alleles of all effect sizes are equally as likely to invade provided inbreeding depression is lower than a threshold value. Finally, comparing across cost functions, small effect mutations, are more successful at invading over a wider range of inbreeding depression values under an accelerating or linear curve than under a saturating one.

In conclusion, our models illustrate how pollinators can interfere with mating system evolution, even when the genetic conditions are expected to pave the way for the evolution of complete selfing. By linking attractiveness to the selfing rate, we further provide a mechanism by which pollen discounting can occur with autonomous selfing and consider attractiveness as a cost of outcrossing. In this way, complete selfing can evolve in the face of pollen discounting so long as there is a cost to attraction, but it is still restrictive. The economics of floral investment are not traditionally viewed as a cost to outcrossing, creating a disconnect between models of mating system evolution and floral evolution. Our model illustrates the importance of understanding cost functions for attraction and reward to pollinators, and sets the stage for future models melding ecological, genetic and resource costs to explore conditions that permit or restrict the evolution of pure selfing.

